# The impact of a fatty acid synthase gene in regulating a complex multifunctional trait essential for survival and sexual communication

**DOI:** 10.1101/2025.09.01.673505

**Authors:** Weizhao Sun, Clara Clinder, Mohammed Errbii, Lukas Schrader, Juergen Gadau, Jan Buellesbach

## Abstract

The genetic basis of multi-functional traits shaped by both natural and sexual selection remains poorly understood. In insects, cuticular hydrocarbons (CHCs) are an excellent example of such traits, providing protection against different micro-climatic conditions while simultaneously encoding cues predominantly used in sexual signaling. The *fatty acid synthase* (*fas*) gene family has been implied as an important cornerstone in initiating and maintaining CHC functionality, whereas their exact biosynthetic and regulatory mechanisms have remained poorly understood. Here, we characterize a single fatty acid synthase gene (*fas3*) impacting the main CHC functions in the parasitoid wasp model organism *Nasonia vitripennis*. Knockdown of *fas3* significantly decreases wasp survival under desiccation stress while also completely depleting sexual attractiveness of female wasps, where this trait naturally functions as sex pheromone. Transcriptomic analyses revealed that *fas3* regulates other *fas* and CHC-associated genes, as well as key biosynthetic pathway hubs. We also identified striking sex-specific expression differences in *fas3* across individual developmental stages, suggesting divergent functional roles of this gene in males and females. These findings largely advance our knowledge on the multi-functionality of *fas* genes in governing survival and sexual signaling and underscore their relevance for future studies on metabolomics, ecological adaptation, and sexual communication.

**Author Summary:** Traits that serve both adaptive and reproductive functions—such as those involved in survival and mating success—are often complex, and their genetic foundations remain poorly understood. In insects, chemical compounds on the outer cuticle represent a prime example: they help prevent desiccation and act as key sexual signals. In this study, we investigate the gene *fas3*, a member of the fatty acid synthase family, in the parasitoid wasp *Nasonia vitripennis*, whose knockdown drastically reduces these surface chemicals. Consequently, both of their natural functionalities, namely enhancement of survival in dry conditions and female sexual attractiveness, are severely impaired. Transcriptomic analysis further revealed that *fas3* also regulates multiple other genes, including those in further major biosynthetic pathways. Our findings shed light on how a single gene can coordinate the expression of multifunctional traits, contributing to both ecological adaptation and sexual communication.

## Introduction

Understanding the genetic basis of complex traits that serve multiple biological functions remains a central challenge in evolutionary biology and genetics. Traits that mediate both local adaptations to environmental conditions while simultaneously affecting reproductive success have been hypothesized to evolve under divergent selection pressures due to their distinct functional roles (Servedio et al. 2011; Haller et al. 2012; Sakamoto and Innan 2020). For instance, the divergence of beak morphologies in the Darwin finch genus *Geospiza* affects both their foraging niches as well as their mate choice (Podos et al. 2004; Huber et al. 2007). Similarly, coloration in *Heliconius* butterflies and *Hypoplectrus* reef fishes serve as both anti-predatory adaptation and sexual signals (Puebla et al. 2007; Jiggins 2008; Salis et al. 2019). However, most empirical studies on such multifunctional traits focus on the phenotypic level, while their underlying genetic architecture—critical for understanding heritability and evolutionary potential—remains comparatively underexplored, particularly for traits with high levels of complexity (Servedio et al. 2011; Thibert-Plante and Gavrilets 2013).

Cuticular hydrocarbons (CHCs) in insects are an excellent example of complex, multifunctional traits that remain both phenotypically as well as genetically tractable (Blomquist and Ginzel 2021; Holze et al. 2021). These waxy lipids covering the cuticle of terrestrial insects and other arthropods unite the provision of a protective layer against desiccation as well as the functionality as key chemical cues predominantly used in sexual communication (Blomquist and Bagnères 2010; Chung and Carroll 2015). Their protective role against water-loss renders them pivotal in the adaptation to different micro-climates (Gibbs 1998; Gibbs and Rajpurohit 2010; Buellesbach et al. 2018b), while their predominance in sexual communication is frequently associated with mate preference and assortative mating (Gulias Gomes et al. 2008; Blomquist et al. 2020; Heggeseth et al. 2020). Chemically, CHC profiles can be highly complex, often comprising dozens to hundreds of different long-chained compounds with partially diverse branching patterns (methyl-branched alkanes) and degrees of unsaturation (olefines). CHCs are primarily synthesized and modified in oenocytes, a group of specialized glandular cells beneath the epidermis (Wigglesworth 1933; Ferveur et al. 1997; Schal et al. 1998).

CHC biosynthesis is closely linked to fatty acid metabolism and governed by an intricate enzymatic regulatory network (Liu and Li 2020; Holze et al. 2021). Briefly, both processes start with the conversion of acetyl-Coenzyme A (CoA) to malonyl-CoA which is then further extended by fatty acid synthases (FAS) to produce long-chain fatty acids (LCFAs) (Juarez et al. 1992; Gu et al. 1997; Barber et al. 2005). The incorporation of methyl-malonyl-CoA subunits at certain positions generates different methyl-branching patterns whereas desaturases introduce double bonds (Cook 1985; Nelson and Blomquist 1995). Subsequently, LCFAs are elongated to produce very long chain fatty acids (VLCFAs), which then undergo reduction, dehydrogenation and decarboxylation, to yield the final CHC products (Howard and Blomquist 2005; Holze et al. 2021).

The earliest insights into the genetic basis of CHC biosynthesis came from studies on *Drosophila*, which identified several major genetic loci associated with sex-and species-specific CHC variation (e.g. Ferveur 1991; Coyne and Oyama 1995; Ferveur and Jallon 1996). However, the subsequent, almost exclusive focus on this model organism for studies on CHC genetics restricted a phylogenetically broader view on the genetic factors responsible for maintaining CHC functionality across wider taxonomic lineages (Ferveur 2005; Mallet 2006; Holze et al. 2021). Yet, more recently, candidate genes have been found in a wider array of insect taxa, allowing for a more comprehensive picture on the genetic protagonists governing CHC functionality (Dembeck et al. 2015; Balabanidou et al. 2016; Sun et al. 2023). Particularly the *fatty acid synthase* (*fas)* gene family has been shown to impact CHC composition and variation in several insect taxa (Wicker-Thomas et al. 2015; Holze et al. 2021). Multiple characterized *fas* genes in the German cockroach *Blattella germanica (*Pei et al. 2019), the migratory locust *Locusta migratoria* (Yang et al. 2020), and the kissing bug *Rhodnius prolixus* (Moriconi et al. 2019) affect CHC profile compositions in various ways as well as physiological properties mostly related to desiccation survival. In one particular case, the knockdown of a *fas* gene *(FASN2*) also decreased male mating success in *Drosophila serrata* (Chung et al. 2014). However, the regulatory hierarchy among *fas* genes within the larger CHC biosynthesis framework have remained largely uncharacterized. It is also unclear how *fas* gene expression contributes to maintaining the multifunctionality of CHCs in both environmental adaptation and sexual signaling.

The present study attempts to close these knowledge gaps by characterizing the function of a specific *fas* gene, *fas3*, in the parasitoid wasp *Nasonia vitripennis,* a model organism for the economically and ecologically important insect order Hymenoptera (Gadau et al. 2008; Werren et al. 2010; Li et al. 2023). We first investigate how *fas3* knockdown affects CHC profiles and their associated biological functions. Silencing *fas3* dramatically reduces overall CHC quantities in both male and female wasps and significantly lowers desiccation survival rates in dry conditions in both sexes. Moreover, it completely eliminates sexual attractiveness of female wasps, where CHCs naturally function as sex pheromones (Steiner et al. 2006; Buellesbach et al. 2013). Further investigations revealed strong sex-specific differences in *fas3* expression across developmental stages, indicating distinct functional roles in males and females. Finally, comprehensive gene co-expression network analyses reveal that *fas3* regulates other *fas* and CHC-related genes, also revealing potential upstream regulators and connections to the general fatty acid metabolism. Together, these findings challenge the view of *fas* genes as functionally isolated units and instead support a more integrated, hierarchical role in regulating CHC biosynthesis and its multifunctionality.

## Results

### Knockdown of *fas3* dramatically reduces overall CHC quantities

We performed dsRNAi knockdowns of *fas3* in male and female *Nasonia vitripennis* yellow pupae and conducted gene expression and CHC analyses in the emerged adult wasps. We confirmed the successful knockdown of *fas3* through qPCR, demonstrating that the relative *fas3* expression was significantly downregulated in the RNAi knockdown males (*fas3* RNAi vs WT: p<0.001; *fas3* RNAi vs GFP RNAi: p<0.001) and females (*fas3* RNAi vs WT: p<0.001; *fas3* RNAi vs GFP RNAi: p<0.01) compared to control wasps (Fig. 1 A&D). Subsequently, we analyzed the CHC profiles of *fas3* RNAi wasps, based on 54 identified and quantified individual CHC compounds (Fig. 1 B&E, Table S1). Knockdown of *fas3* led to a dramatic decrease in overall CHC quantities in both sexes, compared to that of the controls (p<0.001 in each comparison, Fig. 1 C&F). In both males and females, methyl-branched alkane quantities are largely reduced compared to both control groups (p<0.001 in each comparison, Fig. S1), which comprise the vast majority of compounds in *N. vitripennis* CHC profiles (Buellesbach et al. 2022). Furthermore, only females show both a clear reduction in *n*-alkane (p<0.001 vs. both controls) as well as an increase in *n*-alkene (p<0.001 vs. both controls) quantities, although the latter compound class only occurs in comparatively small proportions in *N. vitripennis* females (Buellesbach et al. 2022).

**Figure 1.**
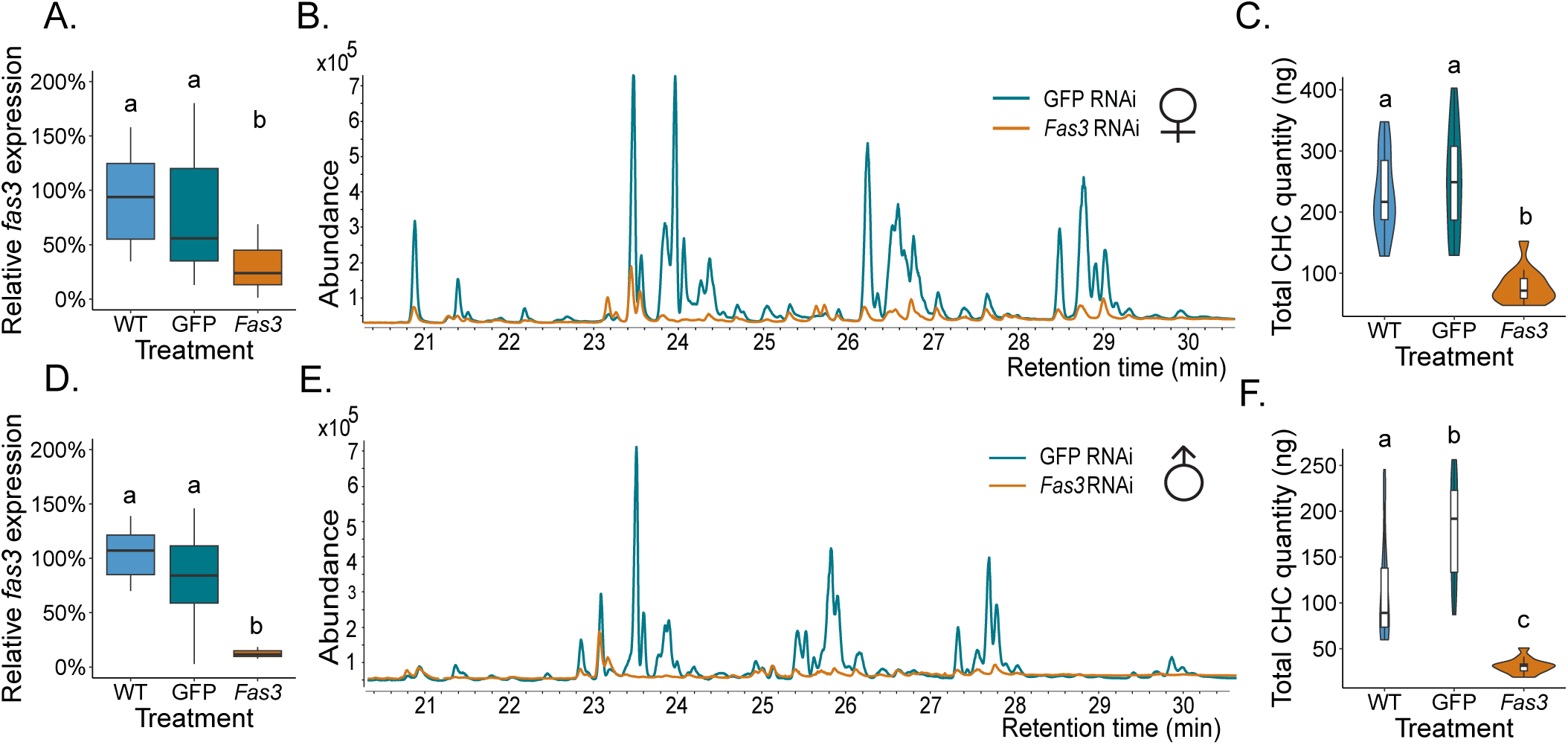
Knockdown of *fas3* significantly reduces overall CHC quantities in *N. vitripennis* males and females. A) Relative *fas3* expression in WT (N = 15), GFP RNAi (N = 16) and *fas3* RNAi (N = 15) females (indicated by blue, green, and orange boxplots, respectively, this color scheme is consistently applied for the following subplots). B.) Comparison of *fas3* (orange) and *GFP* (green) RNAi female FID chromatograms. C.) Total CHC quantities (ng) in WT (N = 14), GFP RNAi (N = 15) and fas3 RNAi (N = 14) females. D.) Relative *fas3* expression in WT, GFP RNAi and fas3 RNAi males (N=15 for each treatment). E.) The comparison of *fas3* and GFP RNAi male FID chromatograms. F.) Total CHC quantities (ng) in WT, GFP RNAi and *fas3* RNAi males (N=15 for each treatment). Significant differences (p<0.05) of relative *fas3* expressions and total CHC quantities between treatments were assessed by Benjamini-Hochberg corrected Mann-Whitney U tests and are indicated by different letters.

### Knockdown of *fas3* decreases desiccation tolerance and depletes female sexual attractiveness

We explored the survival of *fas3* RNAi wasps under different levels of desiccation stress in comparison to both GFP RNAi and WT controls, separately for each sex. In both ambient and low humidity conditions, *fas3* RNAi females showed a significant reduction in survival rates over time in comparison to females of both control groups (p<0.001 in each comparison) (Fig. 2 A-B). *Fas3* RNAi males exhibited significantly lower survival rates (p<0.001) under low humidity conditions but showed no difference in survival rates (p=0.52) in ambient humidity compared to males of both control groups (Fig.2 C-D). We further investigated the mating behavior of unmanipulated WT males on *fas3* RNAi female dummies compared to both GFP RNAi and WT female dummies. This was done to elucidate the impact of *fas3* on the natural function of female CHCs in *N.vitripennis* where they serve as sex pheromones and species-specific cues eliciting male courtship and mating behaviors (Steiner et al. 2006; Buellesbach et al. 2013). Strikingly, not a single male performed mating attempts on *fas3* RNAi female dummies, as opposed to control dummies (p<0.001 vs. both control groups) that elicited natural rates of mating attempts and courtship behavior (70 – 100 %) (Fig2 E; Buellesbach et al. 2013; Sun et al. 2023). To further confirm that the depletion of sexual attractiveness following *fas3* knockdown was due to alterations in CHC profiles rather than other potential cues, we conducted behavioral assays using CHC-cleared *fas3* RNAi dummies reconstituted with freshly extracted CHC profiles from WT females, *fas3* RNAi females, or simply solvent (hexane). Dummies reconstituted with WT female CHC extracts elicited mating attempts from 75% of males, compared to only 42% and 17% for dummies treated with hexane and *fas3* RNAi female CHC extracts, respectively (Fig 2 F). Similarly, dummies reconstituted with WT female CHC extracts elicited courtship from 67% of the tested males, whereas none of the males displayed any courtship towards dummies treated with either hexane or *fas3* RNAi female CHCs (Fig 2 F).

**Figure 2.**
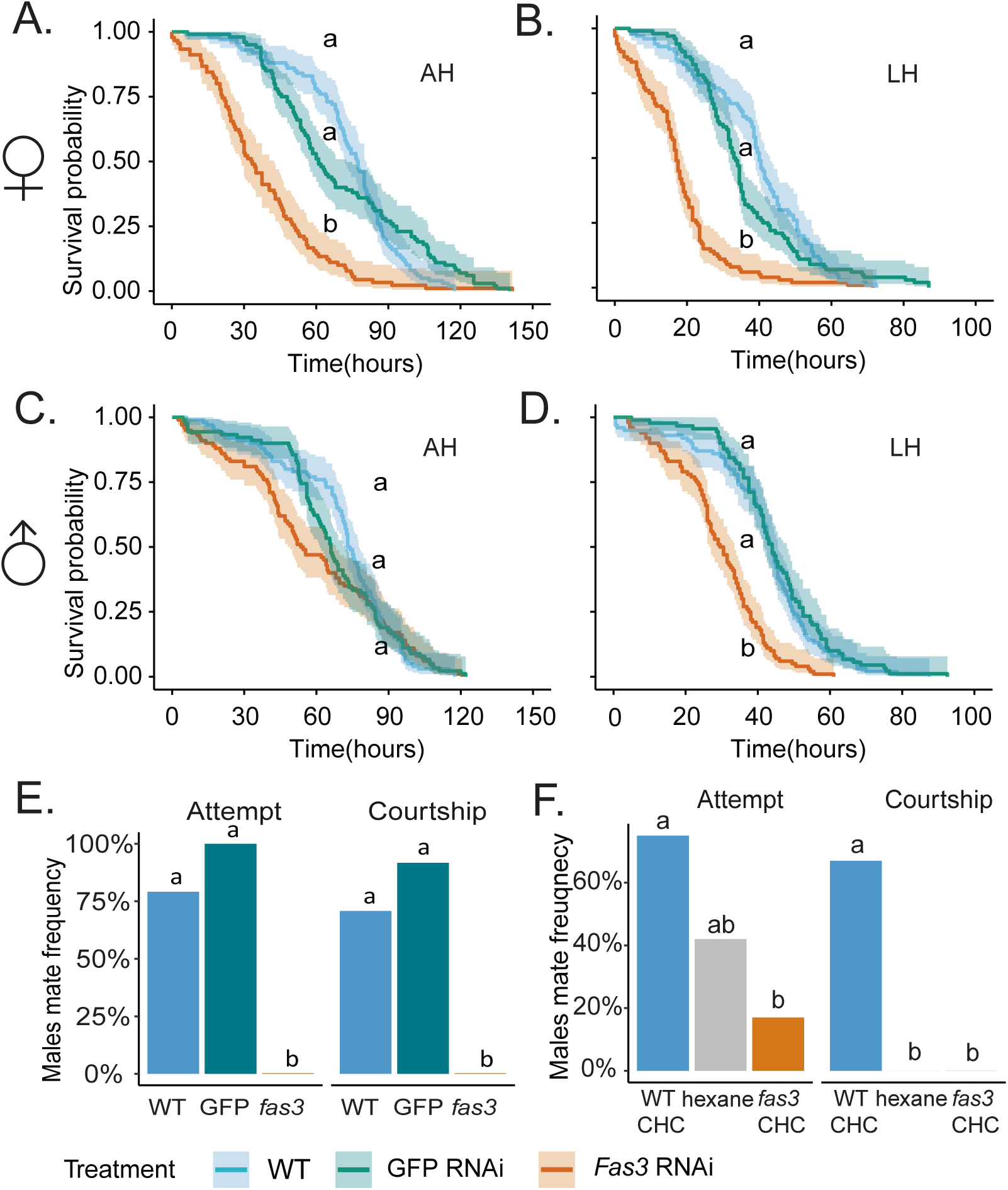
Knockdown of *fas3* decreases survival probability and depletes female sexual attractiveness. A.-B.) Survival probability of WT (blue), GFP RNAi (green) and *fas3* RNAi (orange) females across time under ambient (AH) and low humidity (LH) conditions, respectively, this color scheme is consistently applied for the following subplots. C.-D.) Survival probability of WT, GFP RNAi and *fas3* RNAi males across time under AH and LH conditions. Each treatment group consisted of 10 vials containing 10 wasps each, except for WT males which consisted of 9 vials. Significant differences (p<0.05) of survival probabilities were tested using a Cox regression model that was corrected for multiple testing by the Benjamini-Hochberg procedure and are indicated by different letters. E.) Proportions of males performing mating behavior (attempt and courtship) towards WT, GFP RNAi and *fas3* RNAi freeze-killed females (N=24 for each treatment). F.) Proportions of males performing mating behavior (attempt and courtship) towards CHC-cleared *fas3* RNAi females reconstituted with fresh CHC extracts from WT females, *fas3* RNAi females or hexane alone (N=12 for each treatment). Significant differences (p<0.05) of male mating behavior frequencies towards differently treated females were assessed by Benjamini-Hochberg corrected Fisher’s Exact tests and are indicated by different letters.

### The expression patterns of *fas3* in different developmental stages

To better understand how *fas3* gene expression coincides with CHC profile changes and biosynthesis, we determined the expression of *fas3* across different developmental and adult stages in wild type (WT) *N. vitripennis* wasps. This was done in yellow (yp, 9-10 days after egg deposition), black-and-white (bwp, 11-12 days after egg deposition) and black pupae (bp, 13-14 days after egg deposition) as well as 0, 12 and 24 h old adult wasps after eclosure (A0h, A12h and A24h, respectively) of both sexes. Relative *fas3* expression in male pupae and adults showed a steady increase throughout those six developmental stages (Fig. 3 A). In females, relative *fas3* expression showed a similar increase through the respective pupal stages but began to steadily drop shortly after adult females eclosed (Fig. 3 A). The concordant analysis of CHC profile quantities in different developmental stages showed that generally, *N. vitripennis* pupae exhibit lower overall CHC quantities compared to those of adult wasps (Fig. 3 B-C). However, in adults, overall CHC quantities show a strikingly different trend compared to *fas3* gene expression levels contrasting males and females: Whereas in males, *fas3* gene expression increases largely and steadily through the three adult stages, CHC quantities remain at comparable levels (Fig. 3 C). Contrary to that pattern, in females, despite a steady decrease of *fas3* gene expression from 0-24h of age after eclosure, CHC quantities actually increase during these stages (compare Fig. 3 A and C). A Non-Metric Multidimensional Scaling (NMDS) analysis further showed a clustering of CHC profiles from different pupal developmental stages, clearly distinguishing them from the adult CHC profiles (Fig. 3 D). Moreover, no clear sex-specific differentiation could be detected in the pupal stages, which only begins to manifest in the adults right after eclosure (Fig. 3 D).

**Figure 3.**
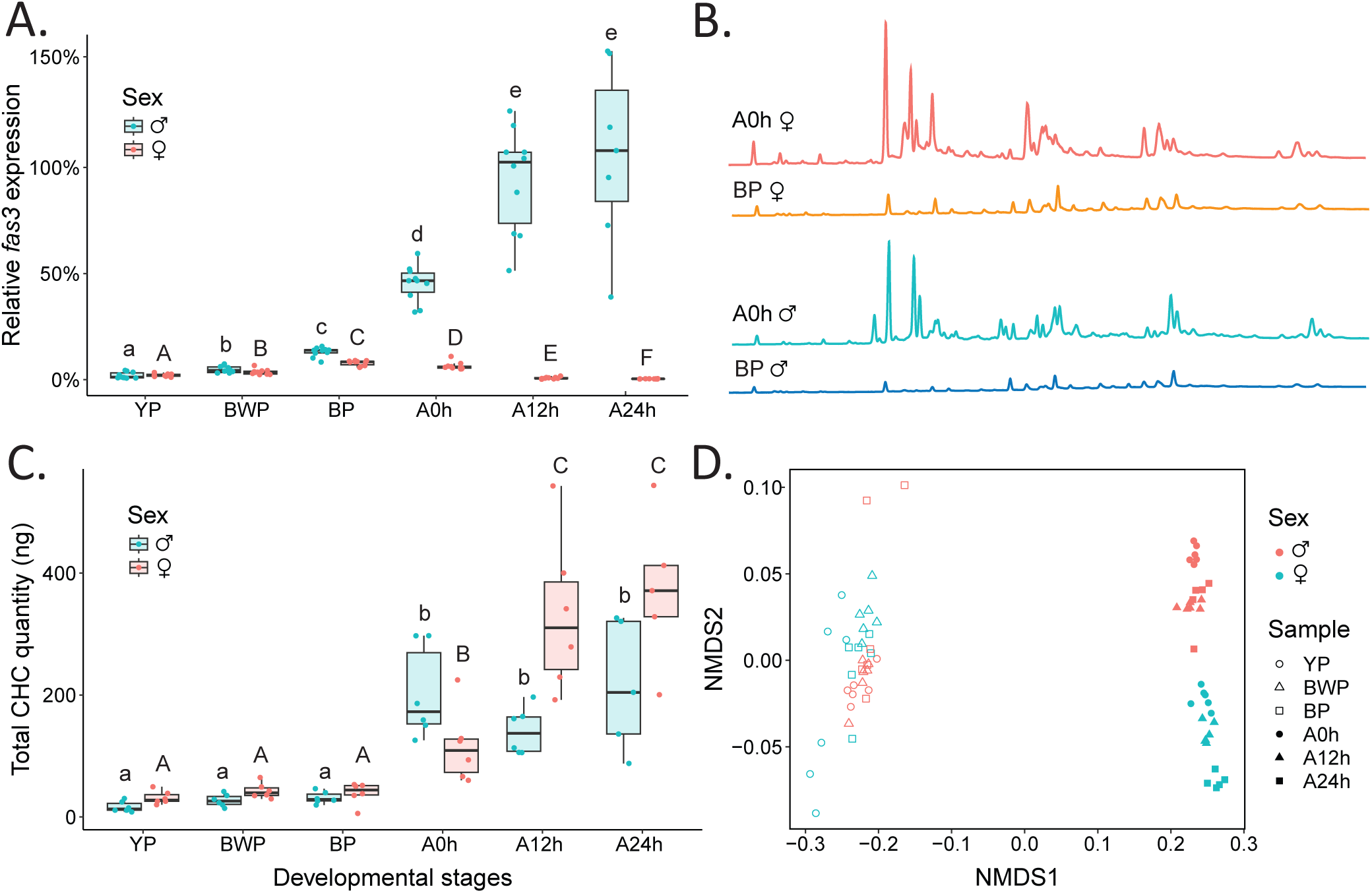
Relative *fas3* expressions and its correlation with cuticular hydrocarbon (CHC) profiles in different developmental stages of *Nasonia vitripennis* A.) Relative *fas3* expression in the pupal developmental stages yellow (yp, 9-10 days after egg deposition), black-and-white (bwp, 11-12 days after egg deposition) and black (bp, 12-13 days after egg deposition) pupae and the adult stages 0, 12 and 24 h after eclosure (A0h, A12h and A24h, respectively). Males are indicated in light blue, females in light red (N=10 for each treatment, except N=7 for Adult24h males). B.) Chromatograms of female and male adults as well as female and male bp males, indicated in orange, dark blue, yellow, and sky blue, respectively. C.) Boxplots depicting the total CHC quantity in the different pupal and adult developmental stages in both sexes as in A), with males in light blue and females in light red (N=6 for each treatment, except N=5 for Adult24h). D.) A Non-Metric Multidimensional Scaling (NMDS) plot illustrating the divergence of CHC profiles between developmental stages (yp in purple, bwp in blue, bp in green, and adults in red) and sexes (females in circles, males in triangles). Significant differences (p<0.05) between gene expression levels (A) and CHC quantities (C) were assessed with Benjamini-Hochberg corrected log-rank tests and are indicated by different letters.

### Differentially expressed genes in *fas3* RNAi females and *fas3* associated gene regulatory network

To gain insights into the gene regulatory network associated with *fas3*, we further sequenced the transcriptomes of *fas3* RNAi, GFP RNAi and WT female abdomens. We restricted ourselves to females in accordance with our main focus on the multi-functionality of CHC profiles, desiccation protection and sexual signaling, the latter of which has only been unambiguously shown in *N. vitripennis* females. A differential gene expression (DEG) analysis revealed three other genes to be significantly down-regulated along with *fas3* in the RNAi knockdown females compared to WT and GFP controls, *fas2* (p<0.001 vs. both control groups), *fas4* (p<0.001 vs. both control groups) and *far18* (p<0.01 vs. both control groups). These results for down-regulated genes remained consistent irrespective of whether the DEG was performed on all annotated genes in the *N. vitripennis* Nvit 2.0 reference genome (Fig. S2 A-B), or on a manually curated list of 190 genes specifically selected for their function in either CHC biosynthesis or fatty acid metabolism (Table S3, Fig. 4 A). However, only in the latter analysis, one gene was found to be significantly upregulated (p<0.05 vs. both control groups), *i.e.,* UPP1 (Fig. 4 A).

**Figure 4.**
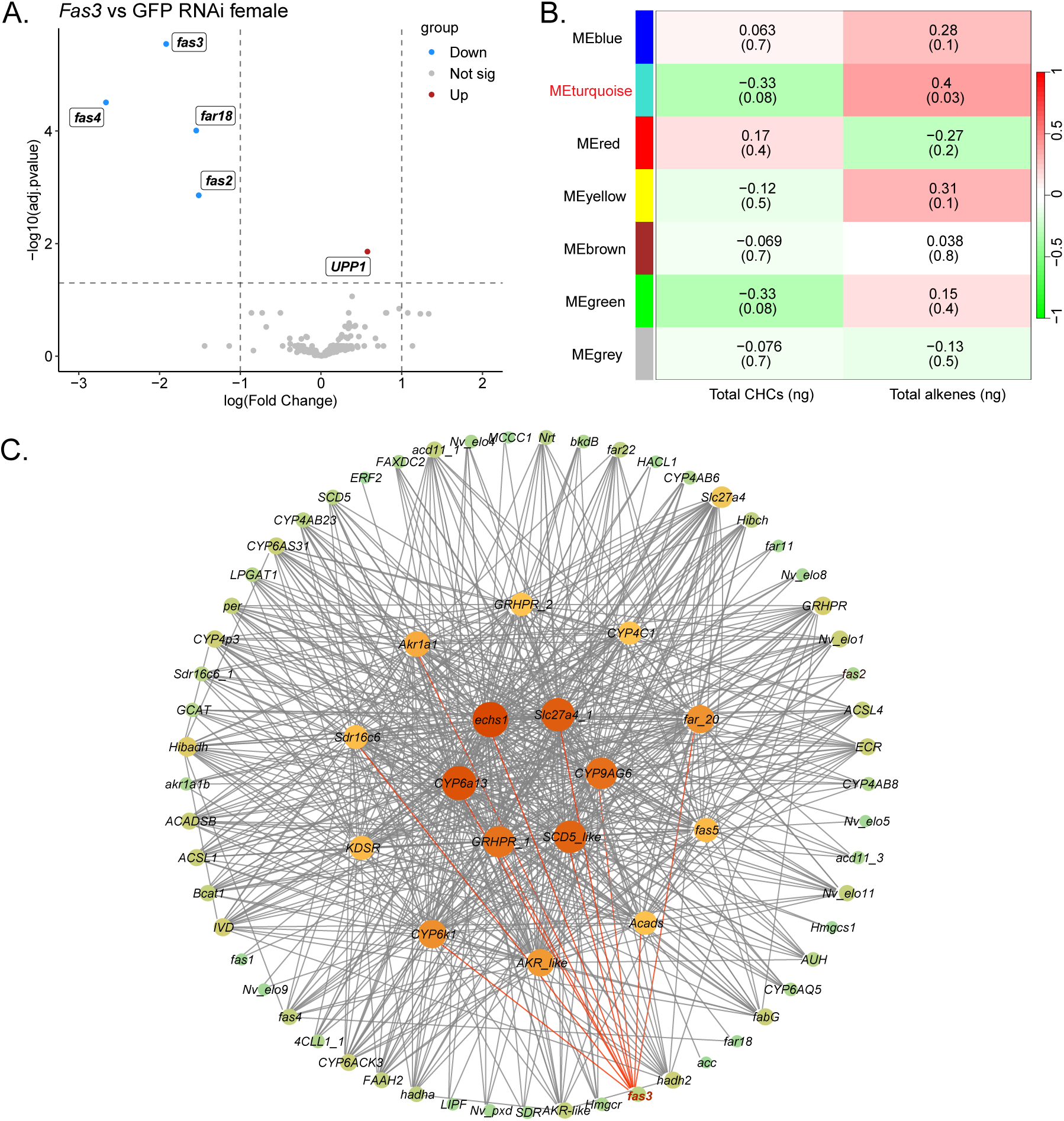
**Differentially Expressed Gene (DEG) analysis and Weighted Gene Correlation Network Analysis (WGCNA) performed with CHC biosynthesis-and fatty acid metabolism-related genes**. A.) A volcano plot illustrating DEGs in *fas3* RNAi (N = 12) females compared to GFP RNAi (N = 11) females. Genes with significant down-regulation are marked in sky blue; all others (no significant changes) are marked in grey. B.) Genes were clustered into modules, indicated by different colors as noted in the module names. Correlations between modules and phenotypic traits (total CHC and *n*-alkene quantities) are represented by colors ranging from green to red (-1 to 1). The correlation coefficients and p-values (in brackets) are indicated for each module. C.) The non-directional gene regulatory network from Module’MEturquoise’ is visualized, with nodes representing genes and grey lines indicating connections (Pearson correlation coefficient > 0.3). The size of a node represents the number of connections a gene has. The node of *fas3* and its connections to other nodes are marked in red.

Proceeding with the selected gene list based on the two metabolic processes most associated with the *fas* gene family (Liu and Li 2020), a subsequently performed Weighted Gene Correlation Network Analysis (WGCNA) clustered genes into seven modules (Fig. S3), which were then correlated with total CHC and total *n*-alkene quantities (Fig. 4 B). Those were chosen based on the inverse effects we uncovered for *fas3* knockdowns on total CHC (down-regulation) and *n*-alkene (up-regulation) quantities (see above and Fig. S1). The module “MEturquoise” exhibited a strong negative correlative trend with total CHC quantities (p=0.08), and a significant positive correlation with total *n*-alkene quantities (p=0.03). For other modules, Pearson correlation coefficients between genes within each module were below a threshold of 0.3 and were thus not further investigated. In the gene regulatory network visualized from the module “MEturquoise” (Fig. 4 C), we identified six hub genes, namely *echs1*, *Slc27a4_1*, *CYP6a13*, *GRHPR_1*, *CYP9AG6* and *SCD5_like*. Notably, *fas3* was found to be directly connected to all six hub genes as well as five additional genes, *CYP6k1*, *Sdr16c6*, *Akr1a1*, *far_20* and *Acads*.

### Expression of all *Nasonia fas* genes in *fas3* RNAi females, *fas* gene phylogeny, in relation to protein domain structures and relative WT *fas* gene

We followed our gene regulatory network investigations with a gene expression analysis to explore the *fas3* knockdown effect on the other identified *fas* genes in the *N. vitripennis* genome, *i.e., fas1, fas2, fas4* and *fas5/6*, the latter constituting a highly similar gene pair most likely originating from a recent tandem duplication (Lammers et al. 2019; Buellesbach et al. 2022; Sun et al. 2023). *Fas4* expression was significantly downregulated in the *fas3* knockdown females (*fas3* RNAi vs WT: p<0.05; *fas3* RNAi vs GFP RNAi: p<0.05) (Fig. 5 A), reflecting similar results as obtained from the transcriptome analysis of *fas3* RNAi females (compare to Fig. 4 A). No significant changes in gene expression were found in the other *fas* genes upon *fas3* knockdown (Fig. 5 A).

**Figure 5.**
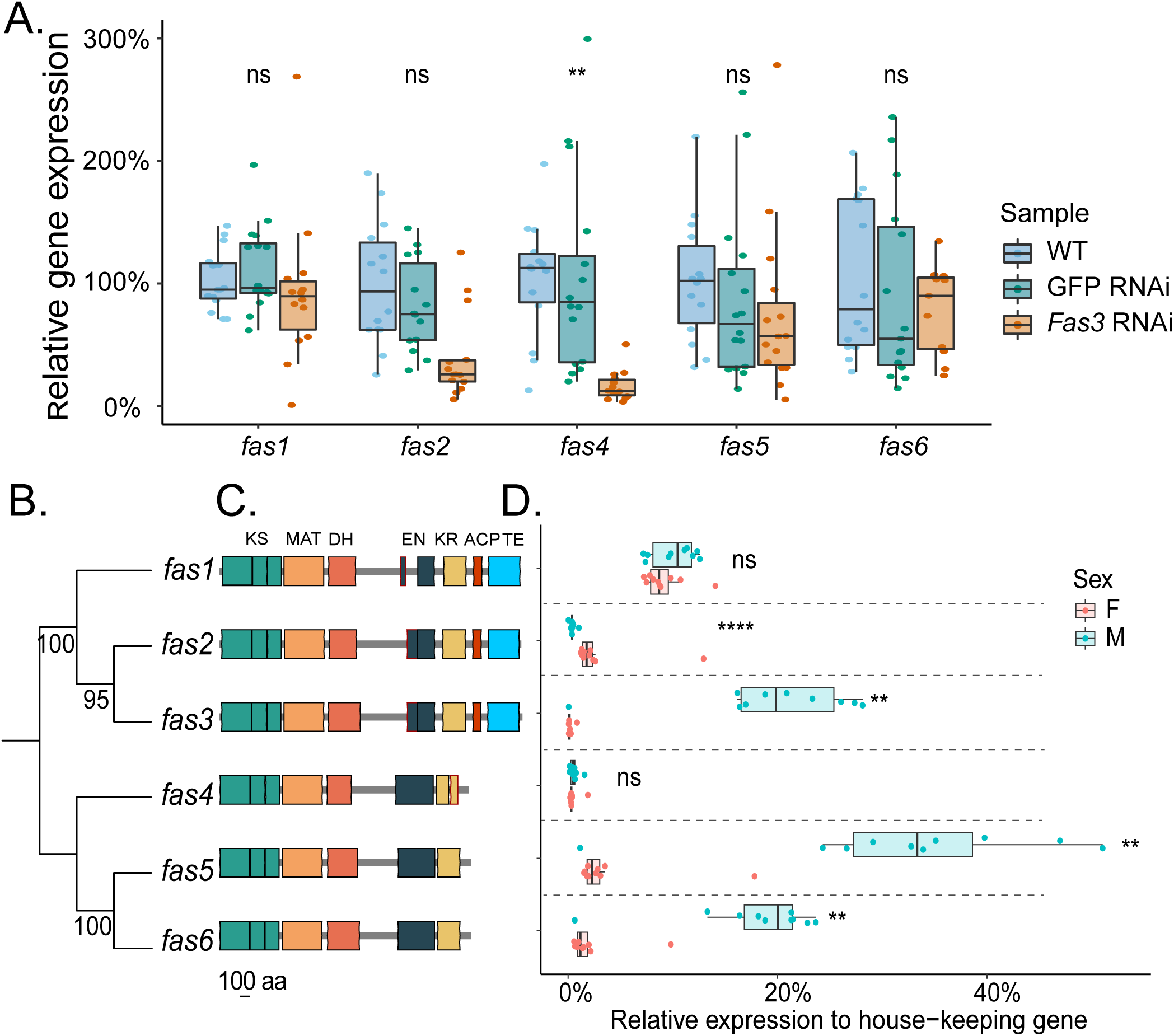
Expression of the other *Nasonia fas* genes in *fas3* RNAi females, *fas* gene phylogeny in relation to protein domain structure, and WT expression patterns of all *fas* genes. A.) Relative gene expressions of the other annotated *fas* genes in *fas3* RNAi females. WT (N = 15), GFP RNAi (N = 16) and *fas3* RNAi (N = 16) treatments were labeled light blue, green and orange, respectively. B.) Phylogenetic tree of the six annotated *fas* genes in *N. vitripennis*. The bootstrap values, based on 100 iterations, are indicated for each branch. C.) Protein domain structures based on the *fas* gene sequences. Seven protein domains are identified and indicated in different colors and acronyms: KS: ketoacyl synthase (in green); MAT: malonyl transferase (in light orange); DH: dehydratase (in dark orange); EN: enoyl reductase (in balck); KR: ketoreductase (in yellow); ACP: acyl carrier protein (in red); TE: thioesterase (in blue). D.) Relative expression patterns of *fas* genes in WT *N. vitripennis* males (light blue) and females (light red) compared to house-keeping genes (N=10 for each sex). Significant differences were assessed by Benjamini-Hochberg corrected Mann-Whitney U tests and are indicated as ns: not significantly different; **: p<0.01; ****: p<0.001.

We further investigated the gene phylogeny between the six *N. vitripennis fas* genes. Two clades could be clearly differentiated, the first containing *fas1*-*3*, and the second, *fas4*-*6* (Fig. 5 B). The phylogenetic relationship of those *fas* genes matched their respective protein domain structures. *Fas1*, *2* and *3* all contain seven domains with similar lengths and orders, *i.e., ketoacyl synthase* (KS), *malonyl transferase* (MAT), *dehydratase* (DH), *enoyl reductase* (EN), *ketoreductase* (KR), *acyl carrier protein* (ACP) and *thioesterase* (TE). Conversely, *fas4*, *5* and *6* only contain the first five domains, while the ACP and TE domains are not present (Fig. 5 C).

Finally, the relative expressions of all six *fas* genes were investigated in WT males and females (Fig. 5 D). Notably, *fas3*, *fas5* & *fas6* showed highly male-biased expression patterns (on average 19.4%, 32.1% and 17.8% in males compared to 0.2%, 3.9% and 2.1% in females, respectively). However, *fas2* showed a low but significantly female-biased expression (on average 2.9% in females, compared to 0.4% in males). *Fas1* has a moderate and non-sex-biased expression (on average 10.1% in males and 9.2% in females). Similarly, *fas4* did not show a sex-biased expression pattern, but a relatively low expression level overall (on average 0.5% in both sexes).

## Discussion

In this study, we characterize a *fatty acid synthase* gene, *fas3*, and demonstrate its central role in shaping a complex, multifunctional trait. RNAi-mediated knockdown of *fas3* led to a dramatical reduction in cuticular hydrocarbon (CHC) quantities, rendering individuals more susceptible to desiccation and simultaneously eliminating the sexual attractiveness of female CHC profiles. These findings substantially advance our surprisingly scarce knowledge on the genetic architecture of complex, multi-functional traits (Servedio et al. 2011; Chung et al. 2014; Chung and Carroll 2015). Our study suggests a rather simple genetic basis required for maintaining the functional threshold in this multimodal trait essential for survival, adaptation, and reproduction with *fas3* emerging as one of the most important mediators for its formation.

Previous work has postulated that insect fatty acid synthases (FAS) generally fall into two broad categories: cytosolic FASs, hypothesized to mainly catalyze the synthesis of precursors to saturated and unsaturated straight-chain CHCs (*n*-alkanes and olefins), and microsomal FASs, speculated to be more specific for methyl-branched CHCs (Juarez et al. 1992; Gu et al. 1997; Holze et al. 2021). However, the functional pattern we observe for *fas3* does not fit neatly into either category. At first glance, the clear downregulation of methyl-branched alkane quantities could be considered consistent with a microsomal role (Fig. S1 B&E). However, the sex-specific impact particularly in females, down-regulating *n*-alkanes while upregulating *n*-alkenes, would more typically indicate a cytosolic FAS function (Fig. S1 A&C).

Interestingly, the recent knockdown of two tandem duplicated *fas* genes (*fas5/6*) in *N. vitripennis* also upregulated *n*-alkene amounts, whereas overall CHC quantities did not change (Sun et al. 2023). Instead, *fas5/6* regulated very specific methyl-branched alkane ratios, strongly arguing for their functionality more downstream of *fas3* and hinting at a hitherto uncharacterized hierarchical regulation among *fas* gene family members (Holze et al. 2021). Comparable complexity has been observed in more recent studies in other insect taxa on *fas* genes. In the German cockroach *Blattella germanica* and the migratory locust *Locusta migratoria*, several *fas* genes have been characterized that either decrease or increase both methyl-branched and straight-chain alkanes (Pei et al. 2019; Yang et al. 2020). Furthermore, two *fas* orthologs, one in *Drosophila melanogaster* (*FASN2*) and one in the kissing bug *Rhodnius prolixus* (*RPRC000123*), affect the down-regulation of methyl-branched alkanes while simultaneously up-regulating straight-chain alkenes (Chung et al. 2014; Wicker-Thomas et al. 2015; Moriconi et al. 2019). Together, these and our findings strongly argue for an extension of the preceding view where *fas* genes are viewed not as specialists for single compound classes but as part of a complementary and potentially hierarchical network for the regulation of CHC variation as a whole.

Our study is also the first to demonstrate a direct functional link between CHC reduction and desiccation sensitivity in *Nasonia*, though this relationship has been demonstrated before in other insect taxa (e.g., Menzel et al. 2019; Wang et al. 2022b; Whyte et al. 2023). Under low humidity, both male and female *fas3* knockdown wasps show significantly reduced survival, strongly suggesting CHC depletion as the primary driver. Interestingly, under ambient humidity, *fas3* knockdown only reduces survival rates in females and not males (Fig. 2 A&C). One straightforward interpretation of this initially surprising finding is that males, which are flightless and remain near their emergence site, may invest differently in water retention mechanisms. Females, in contrast, disperse over larger distances via flight to locate hosts (van den Assem et al. 1980; Grillenberger et al. 2009), potentially trading desiccation resistance for mobility or reproductive flexibility. Furthermore, *fas* gene activity is also closely linked to fatty acid metabolism and therefore, energy storage (Maier et al. 2010; Murphy 2012). Hence, we hypothesize that *fas3* may influence additional sex-specific physiological processes, warranting further investigation into internal lipid composition using more sensitive extraction methods (Flaven-Pouchon et al. 2016), as we are solely focusing on the quantification of external CHCs and their direct impact.

In line with our hypothesis, we observed that native *fas3* expression diverges sharply between sexes following eclosion. While *fas3* expression increases steadily in both sexes during pupal development, adult males maintain high expression levels, whereas females show a steep decline (Fig. 3 A). Paradoxically, CHC quantities rise in females post-eclosion despite the transcriptional *fas3* downregulation (Fig. 3C). This finding strongly suggests that the peak activity of this gene necessary for general CHC production lies in the pupal (and/or very early adulthood) stages for both sexes. Whereas in females gene expression steadily decreases after these stages, the consistent increase in males strongly argues for additional, male-specific *fas3* functionality. For instance, spermatogenesis has been shown to be connected to fatty acid metabolism in other insect taxa and remains yet to be investigated in *Nasonia* (Ben-David et al. 2015; Ng et al. 2015; Kaczmarek and Boguś 2021). Moreover, *N. vitripennis* males continuously emit a long-range pheromone blend consisting of two lactones and 4-methylquinazoline after eclosion (Ruther et al. 2007). Since the biosynthesis of these pheromonal compounds appears to be closely linked to fatty acid metabolism (Ruther et al. 2021), the sustained male-specific expression of *fas3* might be related to their continuous production. These findings open avenues for further research into *fas3*’s possible multifunctionality, especially in males. In females, *fas3* expression peaks during pupal sclerotization and right after eclosion, strongly arguing for this developmental window to be the most crucial for the impact of *fas3* on CHC biosynthesis. Apparently, at least in females, the continuous expression of *fas3* is not necessary for maintaining or augmenting the adult CHC layer, but its silencing during the critical CHC production period cannot be counterbalanced in the adult phase through any other potential CHC maintenance factors (compare Fig. 3 A-C). In addition, sex-specific differentiation of CHC profiles which is particularly crucial for the female-exclusive sex-pheromonal function is only apparent right after eclosion as well (Buellesbach et al. 2013; Mair et al. 2017; Buellesbach et al. 2018a), further corroborating that *fas3* plays no part anymore in CHC biosynthesis after this (Fig. 3 B & D). Concordantly, the *fas3* knockdown renders female CHC profiles completely unattractive to conspecific males, confirming *fas3* as an early upstream key factor in CHC biosynthesis to maintain the threshold for this crucial CHC function as well. This demonstrates how a single gene can govern both physiological integrity and reproductive success.

Furthermore, our transcriptomic comparison revealed only a small set of genes directly affected by *fas3* knockdown, reinforcing its main role as a central catalyst for the biosynthesis of the entire CHC profile rather than a gene regulatory key player (Fig. 4 A). Of the three down-regulated genes in concert with *fas3*, two encompass other *fas* genes (*fas2* and *fas4*) while the other one is a member of the rapidly evolving fatty acyl-CoA reductase (FAR) gene family, *far18* (Finet et al. 2019). The knockdown of *fas2* and *fas4* alone had neither any direct, measurable effects on CHC profiles (Fig. S4), nor on fatty acid metabolism (Lammers et al., 2019), reinforcing their subordinate role. The diverse functionalities of the FAR gene family include catalyzing a later step in CHC biosynthesis (Holze et al. 2021) as well as roles in tracheal gas filling, fertility, and wing expansion (Jaspers et al. 2014; Li et al. 2020), which largely complicates pinpointing a specific effect of this gene.

Strikingly, the only significantly upregulated gene following *fas3* knockdown is an ortholog to UPP1 (uridine phosphorylase 1), an instrumental gene for uridine catabolism, which in turn produces β-alanine and acetyl-CoA (Le et al. 2013; Urasaki et al. 2014). The latter constitutes the initial and rate limiting substrate for CHC biosynthesis, most commonly derived from the citric acid cycle (Barber et al. 2005; Holze et al. 2021). Thus, it makes sense that upon *fas3* downregulation, a key gene from a compensatory, alternative metabolomic pathway gets upregulated that increases the quantity of this crucial substrate for CHC biosynthesis. However, as we have established, this upregulation alone does not rescue the completely downregulated CHC phenotype and appears therefore insufficient to counteract the substantial CHC decrease.

Our gene co-expression network analysis further broadens the scope on *fas3*’s general connections with other metabolic pathways (Fig. 4 C). Notably, *echs1*, encoding the short chain enoyl-CoA hydratase, is critical for catalyzing fatty acid β-oxidation in mitochondria (Janßen et al. 1997). Mutating the *echs1* gene in *Drosophila* flies leads to an immediate shortage of acetyl-CoA supply in the citric acid cycle (Von Ohlen et al. 2012), revealing another connection to this central biosynthetic pathway. Similarly, *GRHPR*_1 encodes a hydroxypyruvate reductase whose up-regulation increases the citric acid cycle rate as well as ATP levels in *Drosophila* (Emtenani et al. 2022). Further genes directly connected to *fas3* appear to be more instrumental for general fatty acid metabolism. *Slc27a4* encodes a long-chain fatty acid transport protein, that contributes to the uptake of LCFAs in all tissues utilizing fatty acids and lipids (Stahl et al. 2001). *Slc* belongs to a family of fatty acid binding proteins that is essential for chitin biosynthesis in the wings of *Drosophila*, whose knockdown even leads to lethal larval phenotypes (Chen et al. 2022). *SCD5* (Stearoyl-CoA Desaturase 5) has been shown to play an essential role the desaturation of fatty acyl-CoA substrates to eventually synthesize monounsaturated fatty acids (Paton and Ntambi 2009). Finally, *CYP6a13* and *CYP9AG6* both encode genes belonging to the Cytochrome P450 Decarbonylase (*CYP4G*) gene subfamily governing vital steps in both CHC and fatty acid biosynthesis (Feyereisen 2020; Liu and Li 2020), uniting these two crucial metabolomic pathways. These uncovered links highlight *fas3* as a nexus of lipid metabolism, integrating CHC biosynthesis with broader biochemical networks.

This intricate link between fatty acid metabolism and CHC biosynthesis through several shared enzymatic pathways (Herman and Zhang 2016; Liu and Li 2020) renders a prediction of *fas* gene functionality in either one very challenging (Fig. 5 B&C). *Fas* genes mainly expressed in the fat body can contribute to the biosynthesis and storage of lipids, which are further transported to different tissues through the hemolymph (Arrese and Soulages 2010; Cheng et al. 2022). For instance, in the cabbage beetle *Colaphellus bowringi*, the gene *FAS2* shows high expression in the fat body and contributes to lipid accumulation, which plays an important role for diapause preparation (Tan et al. 2017). In *Aedes aegypti*, silencing the gene *FAS1* significantly decreases triglyceride and phospholipid levels (Alabaster et al. 2011). Similarly, knocking down *SIFAS1* in *Spodoptera lituna* reduces the content of *a*-lionlenic acid and triglyceride (Song et al. 2022). However, our close investigation of *fas* gene phylogeny as well as FAS protein domain structures defied any clear patterns enabling the prediction of a specific metabolic function for individual *fas* genes according to their sequences, phylogeny or functional domains in our study system (Fig. 5 B-D).

In conclusion, our study revealed the fundamental importance of *fas3* for maintaining a critical phenotypic threshold for CHC functionality with a dramatic impact on both survival and sexual signaling. Furthermore, we unveiled the peak activity and sex-specific differentiation of native *fas3* expression during larval and adult development, hinting at hitherto unexplored further functions governed by this gene. Concordantly, we were able to already demonstrate links of *fas3* with other metabolic networks most commonly associated with lipid synthesis, supporting its status as a major catalyst for several biosynthetic pathways. These findings contribute to our understanding of the genetic architecture underlying multifunctional traits and invite closer examination of fatty acid synthases in the context of metabolomics, ecological adaptation, and sexual communication.

## Materials and Methods

### *Nasonia* strain maintenance and preparation

For all experiments, we used the standard laboratory strain AsymCX for *Nasonia vitripennis*, initially collected from Leiden, the Netherlands. The wasps were reared at a consistent temperature of 25°C, with a relative humidity of 55% and a light-to-dark cycle of 16:8 hours, resulting in an approximate life cycle of 14 days. Pupae of *Calliphora vomitoria* (Diptera: Calliphoridae) were used as hosts.

### RNAi gene knockdown

RNAi Knockdowns were performed following the procedure described in (Sun et al. 2023). Briefly, *fas3* dsRNAs and GFP dsRNA (as a control) were synthesized using MEGA Script T7 kit (Invitrogen, Carlsbad, CA, USA) with the primer pairs listed in Table S2. DsRNA was mixed with 10% red food dye (V2 FOODS, Niedersachsen, Germany) and injected into the abdomens of *N. vtripennis* yellow pupa (9 to 10 days after egg deposition), using a Femtojet microinjector (Eppendorf, Hamburg, Germany). The injected pupae were stored in petri dishes lined with wet tissue for additional humidity under the same conditions as the wasps were reared (see above). Once the pupae emerged as adults, they were collected within 0-24 hours of eclosion and rapidly snap-frozen using liquid nitrogen and stored at-80 °C for further experiments.

### Gene expression analysis

Quantitative PCR (qPCR) was performed to

- assess the efficency of the *fas3* RNAi knockdown by comparing the expression levels of *fas3* (females: N = 15, males: N = 15) with that of GFP knockdown (females: N = 16, males: N = 15) and WT (females: N = 15, males: N = 15) control individuals;
- investigate the expression of *fas3* across different developmental stages (yellow pupa, black-white pupa, black pupa and and 0-24 h old adult wasps), performed separately for males and females (N=10 for each developmental stage, except N=7 for Adult24h males);
- explore the general expression patterns of the six annotated *Nasonia fas* genes in *fas3* RNAi knockdown in females compared to controls (*fas3* RNAi and WT: N = 15, GFP RNAi: N= 16) as well as in WT male and female adult wasps (N=10 for both sexes).

RNA was extracted from each individual wasp using the Quick-RNA Tissue/Insect Kit (Zymo Research, Freiburg, Germany). RNA extraction was immediately performed on the same individuals after their chemical profiles were extracted (see below). Subsequently, RNA was reverse transcribed into complementary DNA (cDNA) using the cDNA Synthesis Kit (CD BioSciences, New York, USA). For the qPCR procedure, we utilized *Nasonia vitripennis* elongation factor 1α (NvEF-1α) as housekeeping gene to measure gene expression levels against, as described in Wang et al. (2022a). The qPCR was performed on a Lightcycler480 qPCR machine (Roche, Basel, Switzerland) with the following cycling conditions: pre-incubation at 95°C for 3 minutes, 40 amplification cycles consisting of 15 seconds at 95°C and 60 seconds at 60°C, and a final step with a standard dissociation curve to verify the amplification specificity.

### Chemical analysis

CHC extraction and analysis procedures followed the protocols outlined in (Sun et al. 2023). Briefly, each individual wasp was immersed in 50 µl of HPLC-grade *n*-hexane (Merck, KGaA, Darmstadt, Germany) in a 2 ml glass vial (Agilent Technologies, Waldbronn, Germany). Vials were then placed on an orbital shaker (IKA KS 130 Basic, Staufen, Germany) and shaken for 10 minutes. Subsequently, the extracts were evaporated under a continuous stream of gaseous carbon dioxide and then reconstituted in a 10 μl hexane solution containing 7.5 ng/μl dodecane (C12) as an internal standard. For analysis, 3 μl of the reconstituted extract were injected in splitless mode using an automatic liquid sampler (PAL RSI 120, CTC Analytics AG, Switzerland) into a gas chromatograph (GC: 7890B) coupled simultaneously to a flame ionization detector (FID: G3440B) and a tandem mass spectrometer (MS/MS: 7010B, all provided by Agilent Technologies, Waldbronn, Germany). The GC system was equipped with a fused silica column (DB-5MS ultra inert; 30 m x 250 μm x 0.25 μm; Agilent J&W GC columns, Santa Clara, CA, USA), and helium was used as carrier gas at a constant flow rate of 1.8 ml/min. The FID operated at a temperature of 300 °C and utilized nitrogen as the make-up gas with a flow rate of 20 mL/min, and hydrogen as the fuel gas with a flow rate of 30 mL/min. The column was split at an auxiliary electronic pressure control (Aux EPC) module, with one portion leading to the FID detector through a deactivated fused silica column piece (0.9 m x 250 μm) at a flow rate of 0.8 mL/min, and the other portion leading to the mass spectrometer through another deactivated fused silica column piece (1.33 m x 250 μm) at a flow rate of 1.33 mL/min. The temperature program for the column started at 60 °C and was held for 1 minute, which was then increased at a rate of 40 °C per minute until reaching 200 °C, followed by a subsequent increase of 5 °C per minute until reaching the final temperature of 320 °C, which was held for 5 minutes.

CHC peak detection, integration, quantification, and identification were performed using the Quantitative Analysis MassHunter Workstation Software (Version B.09.00 / Build 9.0.647.0, Agilent Technologies, Santa Clara, California, USA). CHCs were identified based on their retention indices, diagnostic ions, and mass spectra derived from the total ion count (TIC) chromatograms, whereas their quantification was achieved by simultaneously obtained flame ionization detector (FID) chromatograms, which is the optimal method for hydrocarbon quantification (Agilent Technologies, Waldbronn, Germany, pers. comm.). To determine the absolute quantities of CHCs (in ng), each compound was calibrated using three replicates of a dilution series based on the closest eluting *n*-alkane from a C21-40 standard series (Merck, KGaA, Darmstadt, Germany) with concentrations of 0.5, 1, 2, 5, 10, 20, and 40 ng/µl, respectively. CHC quantities were compared between *fas3* RNAi (females: N = 14, males: N = 15), GFP RNAi (females: N = 15, males: N = 15) and WT controls (females: N = 15, males: N = 15). Furthermore, CHC quantities were compared between different developmental stages (yellow pupa, black-white pupa, black pupa and and 0-24 h old adult wasps, N=6 for each, except for N=5 for Adult24h).

### Behavioral assays

To assess the impact of *fas3* knockdown on female sexual attractiveness, we conducted mating behavior assays by first offering WT, GFP and *fas3* RNAi-treated freeze-killed females (dummies) to 0-48 h old virgin WT males. Second, CHCs were removed from *fas3* RNAi female dummies and reconsitituted with CHC extracts from WT females, *fas3* RNAi females, or with hexane alone as control. CHC removal was achieved by soaking individual female dummies in 50 µL hexane for 3 hours. For reconstitution, freshly extracted female CHC profiles (extraction method as described above) were resuspended in 5 µL hexane and applied to a CHC-cleared dummy. After the complete evaopration of the solvent, the CHC-reconstituted female dummies were offered to 0-48 h old virgin WT males. All behavioral assays were conducted in a mating chamber consisting of two identical aluminum plates (53 x 41 x 5 mm each). Each plate had 12 drilled holes (6 mm diameter) that served as observation sites. To prepare for recording of the behavioral assays, individual female dummies were placed in each hole of one plate, while individual WT males were placed in each hole of the opposite plate and immediately covered with glass slides (Diagonal GmbH &Co. KG, Münster, Germany). To initiate the behavioral assays, the two plates were quickly joined together. The assays were then recorded with a Canon camera (EOS 70D, Tokyo, Japan) for a duration of 5 minutes. All behavioral assays were conducted inside an enclosed wooden box with consistent illumination (100 lumens) provided by an LED light (L0601, IKEA Dioder, Munich, Germany). The males’ behavior towards the females was subsequently assessed based on the presence or absence of two consecutive behavioral displays: mating attempt, which involved physical contact between the male antennae towards the female’s body surface, and courtship behavior, characterized by a series of stereotypic head nods and antennal sweeps after mounting the female (van den Assem et al. 1980). These behaviors have been established as clear indicators of sexual attractiveness of female CHC profiles (Buellesbach et al. 2013; Buellesbach et al. 2018a; Sun et al. 2023). Their individual occurrences were scored during the recordings and compared between WT, GFP RNAi and *fas3* RNAi females (N=24 for each treatment).

### Desiccation assays

Dessication assays were conducted to investigate the impact of *fas3* knockdown on the survival of wasps under different levels of desiccation stress. High desiccation stress conditions were created by introducing 0.6 g desiccant (DRIERITE, Merck, KGaA, Darmstadt, Germany) into transparent plastic vials (76mm height, 10mm diameter), which were airtightened with rubber plugs, leading to low (9%) relative humidity after 24 h. Within each vial, we placed a cotton piece and a stainless-steel grid in the center to separate the desiccant and the remaining space of the vial (∼35 mm height), which served as an observation site where individual wasps were carefully placed throughout the duration of the desiccation assay. For control vials with moderate desiccation stress under ambient conditions (55% rel. humidity), vials were prepared similarly but without adding the desiccant. The relative humidity in the test tubes was monitored using a humidity–temperature probe (Feuchtemesssystem HYTELOG-USB, B+B Thermo-Technik GmbH, Donaueschingen, Germany) with a measurement accuracy of ±2% relative humidity at 25°C. To prepare for the desiccation assays, newly eclosed male and female wasps (0-24 hours old) were collected, sorted into groups of 10, and provided with honey water (Bluetenhonig, dm-drogerie markt GmbH & Co. KG, Karlsruhe, Germany) for 9 hours. Subsequently, each group of wasps was randomly assigned to the previously prepared vials with either high or moderate desiccation stress. Recordings were conducted using a looped VLC media player script (VideoLAN, Paris, France), which initiated a 2-minute recording (Logitech C920 HD PRO webcam, Logitech GmbH, München, Germany) every 2 hours until the last wasp ceased movement on the grid. The number of surviving wasps in each vial was assessed, and based on these numbers, survival curves were constructed and compared among WT, GFP RNAi and *fas3* RNAi treatment groups (*i.e.,* vials with 10 wasps each, N = 10 for each treatment group except N = 9 for WT males). The Cox Proportional Hazards Model with the R package’survival’ was used to assess the survival probability, in which the knockdown treatment was designated as the fixed factor, and variation among replicates was treated as a random factor. Statistical significance was evaluated using the log-rank test (Therneau and Grambsch 2000), and to address multiple testing across different humidity treatments, the Benjamini–Hochberg procedure was applied for correction (Benjamini and Yekutieli 2001).

### RNAseq methodology

RNA was extracted from the abdomens of female wasps (N=11 for WT and GFP RNAi, N=12 for *fas3* RNAi), directly after their CHC profiles had been extracted. Subsequently, we performed library construction for transcriptome sequencing samples using the NEBNext Ultra II Directional RNA Library Prep Kit for Illumina. Paired end sequencing (100 bp) was performed at an Illumina HiSeq 2500 sequencer. Prior to mapping, the quality of the raw reads was assessed using FastQC (v.0.11.7) and Trimmomatic (v.0.38) (Bolger et al. 2014) was used to remove short and low-quality reads as well as adapter sequences (options: ILLUMINACLIP: TruSeq3-PE.fa:2:30:10 SLIDINGWINDOW:4:20 MINLEN:30). Additionally, we filtered out ribosomal RNA (rRNA) sequences using SortMeRNA (Kopylova et al. 2012) and the silva databases (Quast et al. 2012). The resulting rRNA-free paired-end reads were then mapped to the Nvit_psr_1.1 reference genome (accession: GCA_009193385.2) using STAR (STAR_2.6.0c) (Dobin et al. 2012) and mapping quality was evaluated with Qualimap (Okonechnikov et al. 2016). We used FeatureCounts (v2.0.3) (Liao et al. 2013) to count the number of reads per annotated gene in the reference for downstream differential gene expression analysis. On average, 94% of the reads mapped to a unique position in the genome and over 73% of the reads mapped to exons and were used to produce the count data.

### Gene selection for CHC biosynthetic pathway and fatty acid metabolism

A list of 190 genes associated with the CHC biosynthetic pathway and fatty acid (FA) metabolism were manually selected from all annotated genes in the *N. vitripennis* Nvit 2.0 reference genome (Rago et al. 2016) (Table S3). The selected genes included: a) the orthologues of CHC biosynthesis-related candidate genes in *Nasonia* genome, which were identified by Buellesbach et al. (2022); b) genes annotated under the Gene Ontology terms related to fatty acid metabolism; c) genes featuring specific descriptors, *e.g.,* “-CoA”, “carbon chain elongation”, “elongase” and “reductase”. The curated short gene list was subsequently used in the transcriptomic analysis, which enabled us to focus on and narrow down the genetic changes in the CHC and FA biosynthetic process.

### Differentially expressed genes (DEG) analysis

We conducted a differentially expressed genes (DEG) analysis using the R package “limma” (Ritchie et al. 2015). Read counts were converted to log2 counts per million (CPM) and genes were filtered for those with at least 20 reads. We applied the method of trimmed mean of M-values (TMM) (Robinson and Oshlack 2010) from edgeR (Robinson et al. 2010) to calculate normalization factors and to account for library size variations. Differentially Expressed Genes (DEGs) were identified using a significance threshold (p<0.05) with Benjamini-Hochberg correction (Benjamini and Yekutieli 2001). DEG analysis was performed both on all annotated genes in the *N. vitripennis* Nvit 2.0 reference genome (Rago et al. 2016) and the short-listed genes associated with CHC biosynthesis and FA metabolism (see above).

### Weighted Gene Correlation Network Analysis (WGCNA) analysis

We employed the “WGCNA” R package (Langfelder and Horvath 2008) to construct a gene co-expression network associated with the knockdown of *fas3*. Initially, the samples were clustered to identify and remove outliers. Subsequently, the pariwise Pearson correlations between each pair of genes were calculated. We further selected an appropariate soft-thresholding power (β) which fits the scare-free topology criterion. Afterwards, we raised the pairwise correlations to the power to create an adjacency matrix which represents the connection strength between genes. The adjaceny matrix was used to calculate the topological overlap matrix (TOM), based on which a hierarchical clustering was performed to identify modules comprising highly interconnected genes. Within each module, we calculated the module eigengenes to summarize the expression patterns of the genes within. These module eigengenes were further correlated with phenotypic traits, including total quantity of all CHCs and *n*-alkenes. Modules showing significant correlations with these phenotypic traits were further extracted and visualized in the software Cytoscape v3.10.1 (Shannon et al. 2003).

### *Fas* gene phylogenetic tree construction and protein domain comparision

To determine the phylogenetic relationship of all six annotated *fas* genes from the *N. vitripennis* genome (Buellesbach et al. 2022; Sun et al. 2023), their amino acid sequences were obtained from the National Center for Biotechnology information (NCBI) database (Table S2). Then those sequences were aligned in ClustalW to determine sequence similarity and identify conserved regions. Sequences were subsequently trimmed in Gblocks to remove poorly aligned regions. The phylogentic tree construction was performed on the resulting alignment in RAxML (version v8.2.X), using the Maximum Likelihood (ML) method with a bootstrap of 100. The phylogenetic tree was visualized in iTOL (https://itol.embl.de/tree/ 128176106106306041706709767). The protein domain structures of the *fas* gene products were determined and visulized using the webserver CDvist (http://cdvist.joulinelab.org/ static/arc.html?id=230213_falonir&success=true).

## Supplementary data

**Table S1. Comparison of absolute quantities and relative abundances of single CHC compounds.** Indicated are retention indices (RI), CHC compound identifications or possible configurations in case of ambiguities, their mean absolute (ng) amounts with their respective absolute standard deviations (sd) as well as their respective relative amounts (in %) compared between wildtype (WT, females: N = 14, males: N = 15), GFP RNAi (N = 15 for both sexes) and *fas3* RNAi (females: N = 14, males: N = 15) wasps.

**Table S2. List of gene and transcript IDs, primers and the target sequence for the *fas3* knockdown used in the present study.** All primers were designed using the Primer-BLAST tool from the National Center for Biotechnology Information (NCBI). Indicated are the respective gene and primer names, the gene and transcript IDs, the primer sequences, their usage in the experimental protocol, and the target region and sequence of the *fas3* knockdown.

**Table S3. Selected list of 190 genes associated with CHC biosynthesis and fatty acid metabolism.** The list is composed of orthologues to genes verified to impact CHC profiles through targeted knockdown studies (Buellesbach et al. 2022), genes annotated under the Gene Ontology terms related to fatty acid metabolism and genes including descriptors like “-CoA”, “carbon chain elongation”, “elongase” and “reductase”. Indicated are NCBI gene IDs, annotations and descriptions.

**Figure S1. Absolute quantities (ng) of different compound classes in differentially treated male and female wasps** A.-C.) The total absolute *n*-alkane, methyl-branched (MB) alkane and *n*-alkene quantities in WT (blue, N = 14), GFP RNAi (green, N = 15) and *fas3* RNAi (orange, N = 14) females, respectively. D.-F.) The total absolute *n*-alkane, methyl-branched (MB) alkane and *n*-alkene quantities in WT (blue), GFP RNAi (green) and *fas3* RNAi (orange) males, respectively, N=15 for each treatment. Significant differences between treatments were assessed by Benjamini-Hochberg corrected Mann-Whitney U tests.

**Figure S2. Differentially Expressed Gene (DEG) analysis performed on all annotated genes in the *N. vitripennis* reference genome A.)** Volcano plot illustrating DEGs in *fas3* RNAi (N = 12) compared to GFP RNAi (N = 11) females on a log10 scale using a Benjamini-Hochberg corrected significance threshold of p < 0.05, genes with significant down-regulation are marked in sky blue; all other (no significant changes) are marked in grey. B.) Volcano plot illustrating DEGs in *fas3* RNAi (N = 12) compared to GFP RNAi (N = 11) females using a log2 fold change scale, short-listed genes according with functionalities in either CHC biosynthesis or fatty acid metabolism are indicated in red.

**Figure S3. WGCNA analysis soft threshold powers and co-expression tree**. A.) Scale-free topology index and gene mean connectivity were simulated for various soft-threshold powers. B.) Co-expression tree indicating how the selected genes were clustered into the different modules, MEblue, MEturquoise, MEred, MEyellow, MEbrown, MEgreen and MEgrey, compare to Fig. 4.

**Figure S4. Knockdown of *fas2* and *fas4* had no direct impact on CHC profiles of *Nasonia vitripennis*** A.) Relative *fas2* expression between GFP-and *fas2*-RNAi injected wasps B.) Relative *fas4* expression between GFP-and *fas4*-RNAi injected wasps C.) Chromatographic comparison of CHC profiles from adult GFP-and *fas2*-RNAi injected wasps D.) Chromatographic comparison of CHC profiles from adult GFP-and *fas4*-RNAi injected wasps. Statistical comparison for A. and B. performed with Mann-Witney U tests.

## Author Contributions

J.B. conceived the study; J.B. & W.S. designed the study; W.S., & C.L. performed the experiments and collected the data, W.S., C.L. & M.E. analyzed the results; W.S., M.E., L.S. & J.B. validated the results, W.S. &, M.E. & J.B. visualized the data; J.B. & J.G. provided the resources; W.S. & J.B. wrote the original draft; J.B., W.S., L.S. & J.G. reviewed and edited the final manuscript version, J.B. & J.G. supervised the project; J.B. acquired the funding for the study.

## Acknowledgements

We would like to thank Kathrin Brüggemann and Hildegard Schwitte for assisting with the experiments. Furthermore, we thank Sabine Nooten, Barbara Feldmeyer and Jaime Mauricio Anaya-Rojas for helpful suggestions on the first draft of this manuscript.

## Funding

This research was funded through a grant by the Deutsche Forschungsgemeinschaft (DFG, German Research Foundation) – 427879779 to (BU3439/1-1). The conceptualization of the study was inspired and aided by the framework of the DFG SPP “GEvol” (2349, BU 3439/2-1).

## Data availability

The datasets generated or analyzed during this study will be made available at the figshare data repository.

## Competing Interest Statement

The authors declare that the research was conducted in the absence of any commercial or financial relationships that could be construed as a potential conflict of interest.

## Notes

### Competing Interest Statement

The authors have declared no competing interest.

