## Supplementary figures and images for "The impact of a fatty acid synthase gene in regulating a complex multifunctional trait essential for survival and sexual communication"

### Figure S4.pdf

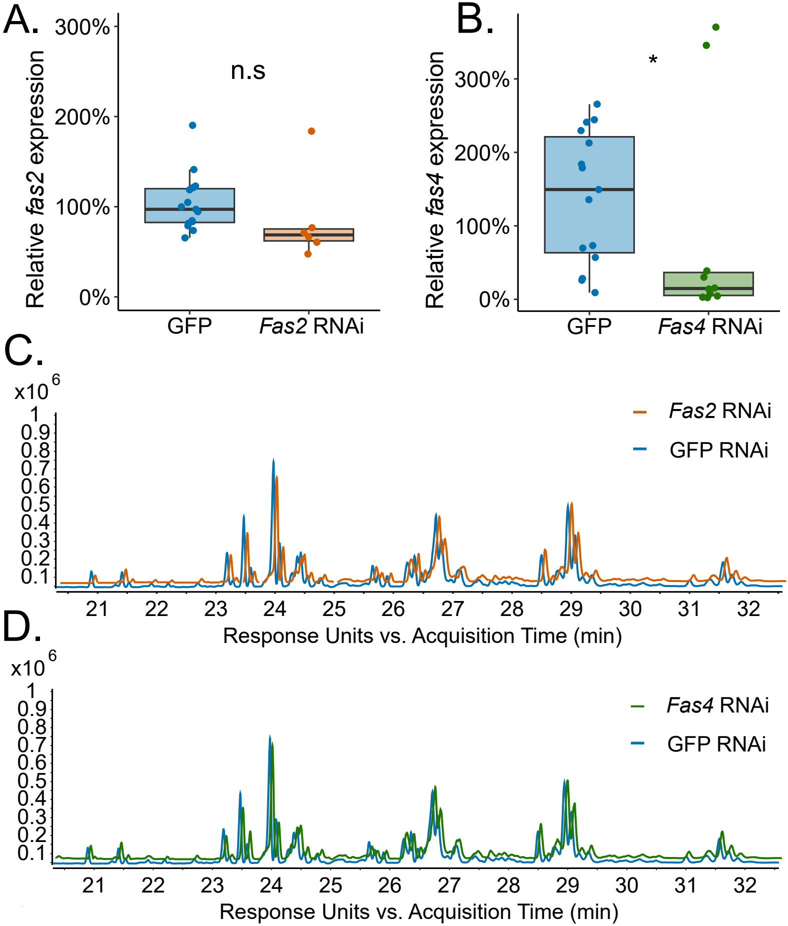

### FigureS1.pdf

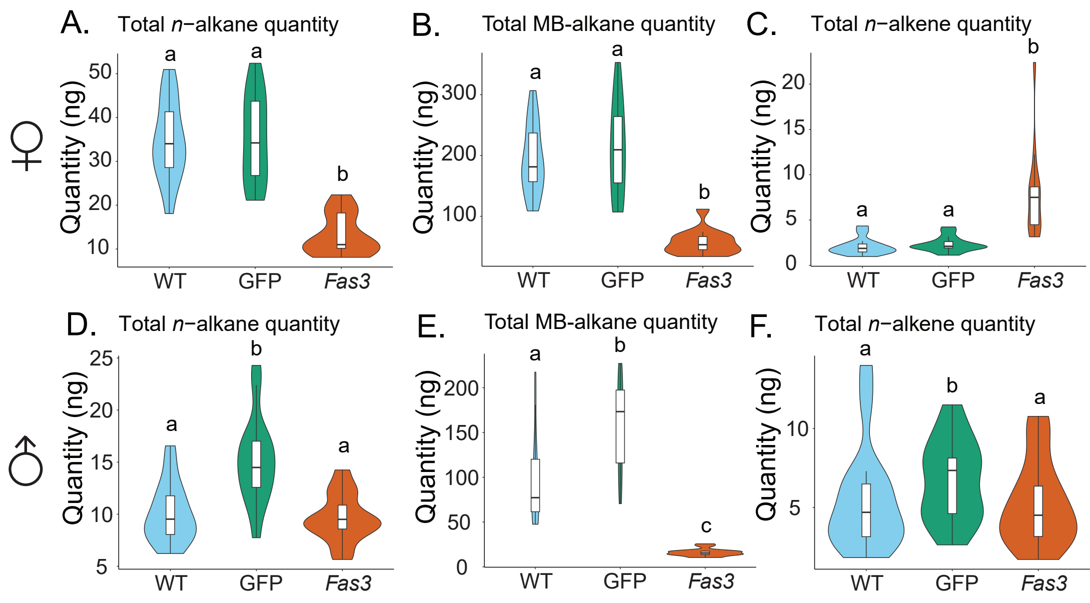

### FigureS2.pdf

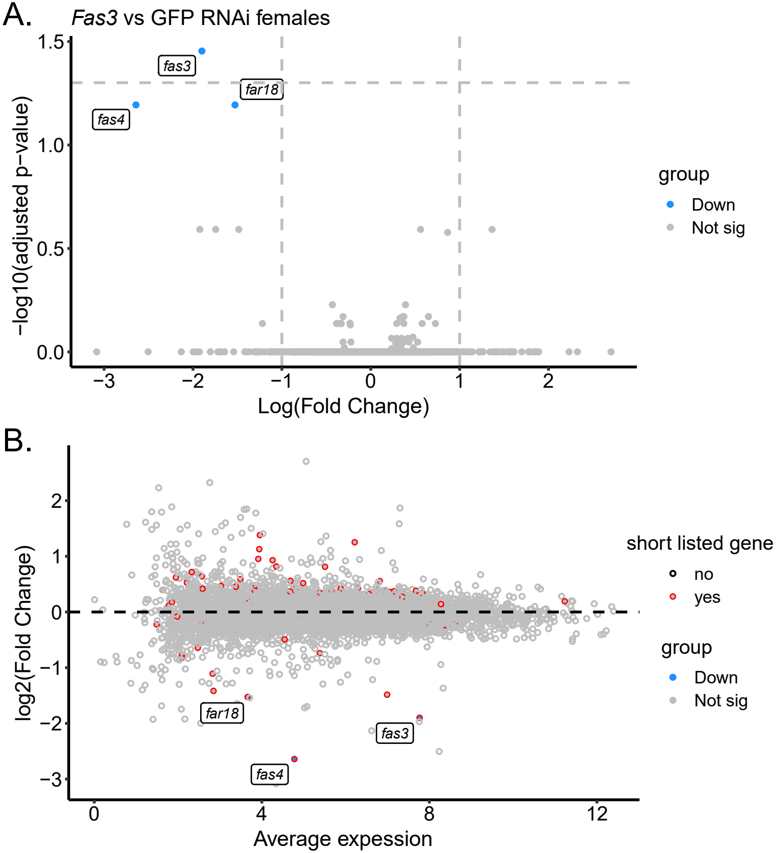

### FigureS3.pdf

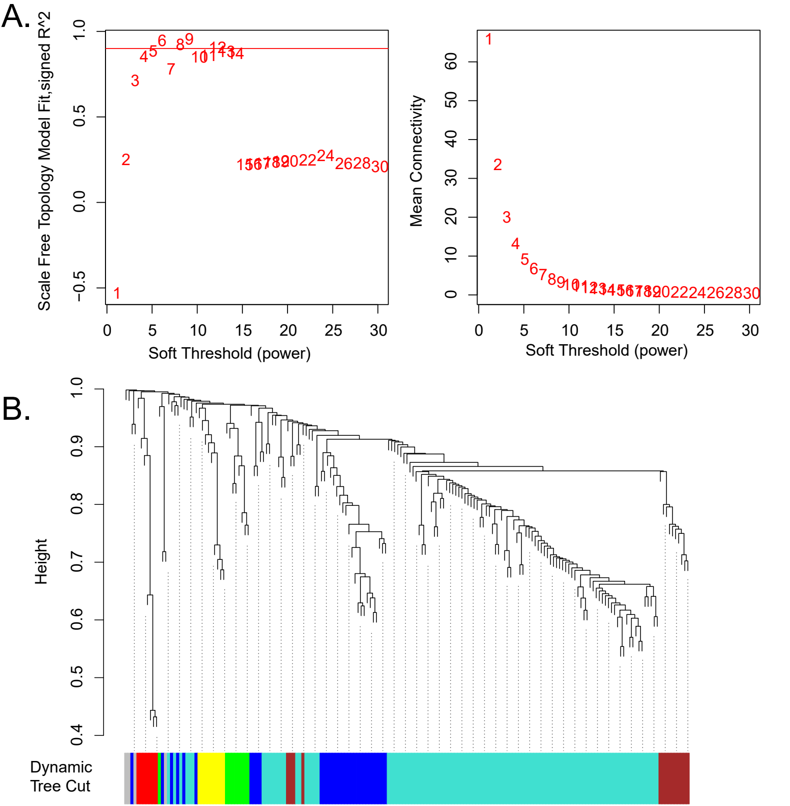
